# Design of readers for pathological PTM proteoforms in Alzheimer’s disease

**DOI:** 10.64898/2026.10.20.760520

**Authors:** Kejia Wu, Tiana Schantz, Lemuel Chang, Prarthna Chabria, Elaina Bashaw, Thejesh Reddy Kalatturu, Doo Yeon Kim, Anqi Chen, Nathaniel Corley, Asim K. Bera, Alex Kang, Hannah Nguyen, Johnny Mendoza, Xinting Li, Magnus S. Bauer, Misa Baum, Brian C. Kraemer, Yuyu Song, David Baker

**Affiliations:** Department of Biochemistry, University of Washington, Seattle, WA, USA; Institute for Protein Design, University of Washington, Seattle, WA, USA; Department of Neurology, Massachusetts General Hospital and Harvard Medical School, Charlestown, MA, USA; Genetics and Aging Research Unit, MassGeneral Institute for Neurodegenerative Disease, Department of Neurology, Massachusetts General Hospital and Harvard Medical School, Charlestown, MA, USA; Geriatrics Research, Education, and Clinical Center, VA Puget Sound Health Care System, Seattle, WA, USA; Division of Gerontology and Geriatric Medicine, Department of Medicine, University of Washington, Seattle, WA, USA; Howard Hughes Medical Institute, University of Washington, Seattle, WA, USA

## Abstract

Alzheimer’s disease (AD) is characterized by heterogeneous Tau phosphorylation and aberrant amyloid-β (Aβ) processing, yet molecular tools that distinguish closely related molecular states of tau and Aβ with predefined specificity remain difficult to develop. Here, we designed de novo protein readers targeting three disease-associated phospho-Tau regions and N-terminal pyroglutamate-modified Aβ (N3pG-Aβ). Screening 192 designs per target identified 51 modification-selective hits among 768 candidates (6.6%). Representative readers exhibited nanomolar to subnanomolar affinities for modified peptides, discriminated authentic modifications from unmodified or substitution-based controls, and preferentially recognized modified full-length targets. We further programmed local modification specificity by interface redesign, generating pT231 readers conditional on the phosphorylation state of neighboring S235 and S237. A 2.0 Å crystal structure of an N3pG-Aβ reader closely matched the computational design model (interface Cα RMSD = 0.52 Å), and redesign of pT231 readers tuned apparent binding affinities from greater than 1 μM to below 1 nM. In a human 3D neuronal culture with both pathological Aβ and tau, selective readers targeting N3pG-Aβ, Tau pT231, and Tau pT217 recognized disease-associated signals, exhibiting colocalization with related antibodies and much higher signal in the AD line than the control. Tau pT231 readers further quantitatively distinguish AD patients and control human brain lysates in dot-blot assays. Together, these results establish genetically encoded readers with programmable specificity for defined Tau and Aβ modification states, enabling molecular-state-resolved detection in human disease models and biological samples, and lay a foundation for controlled pathological molecular perturbation.

## Introduction

Alzheimer’s disease (AD) involves aberrant processing and accumulation of Tau and amyloid-β (Aβ) with underappreciated heterogeneity, yet specific states contributing to neurodegeneration remain difficult to distinguish. Tau undergoes extensive post-translational modifications (PTMs), particularly phosphorylation, that vary across patients and disease stages [1–3] and influence molecular properties and fibril structures [4–6]. Single-molecule measurements reveal nonrandom combinations of phosphorylation sites [7], while specific patterns alter microtubule binding and filament assembly [8–10]. Phospho-Tau species, particularly pT217 and pT231, are important AD biomarkers [11–14], and detecting multiple phosphorylation sites on individual molecules can improve diagnostic discrimination [17]. Aβ is similarly heterogeneous: N-terminal truncation and cyclization generate N3pG-Aβ, a plaque-associated species with distinct aggregation properties [18–20] targeted by donanemab [21–22]. Although phospho-specific antibodies have transformed AD research, their dependence on neighboring modifications is often discovered retrospectively rather than specified during development. For example, AT8 binds triply phosphorylated pS202/pT205/pS208 Tau approximately 30-fold more tightly than pS202/pT205 alone [23–24], while neighboring phosphorylation can enhance or abolish binding by other Tau antibodies [25–28]. Thus, combinatorial PTM specificity often emerges through iterative characterization and remains difficult to predict or control. This complicates measurements of heterogeneous molecular populations sharing nominal modification sites (Fig. 1A). Conventional antibodies face intracellular delivery challenges [30]. While antibody-mediated depletion, Tau-directed degraders, and lysosome-targeted clearance enable protein removal [31–34], compact, genetically encoded readers with predefined modification specificity could distinguish, measure, and ultimately perturb disease-associated states (Fig. 1B). Advances in de novo protein design have enabled recognition of folded proteins [35–36], generative scaffold construction [37–38], and sequence optimization [39], extending to intrinsically disordered regions [40–41] and phosphotyrosine epitopes [42]. RFdiffusion4 extends these capabilities to phosphoserine- and phosphothreonine-containing histone peptides through yeast-display screening of thousands of designs per target, yielding apparent surface affinities of ∼0.14–1 μM [44]. Yet display-derived affinities may not translate to purified proteins, and prospectively specifying recognition of precise combinations of neighboring PTMs remains an unsolved challenge.

**Figure 1.**
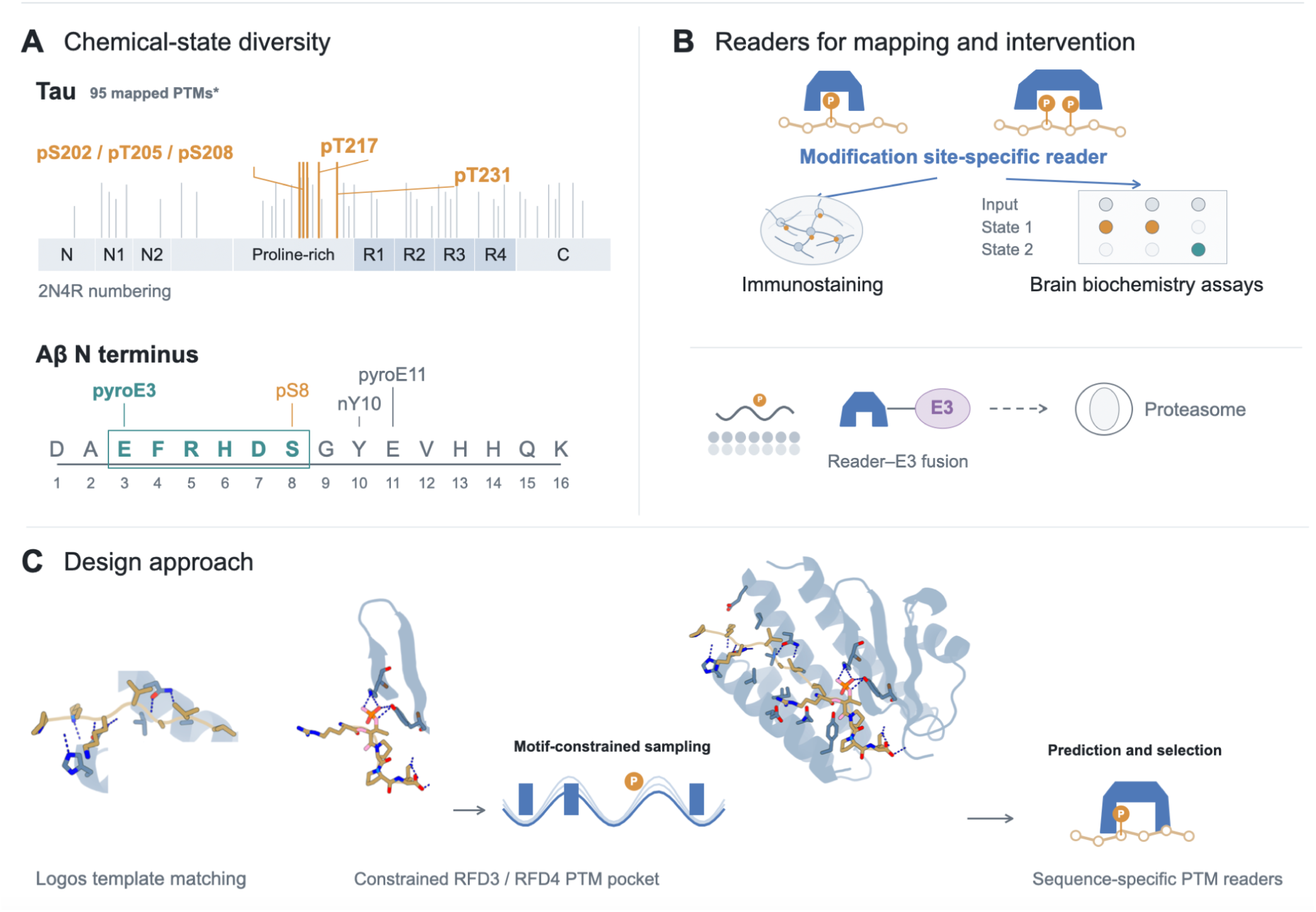
Programmable recognition of locally defined Tau and Aβ modification states. (A) Tau and Aβ modification landscapes motivating reader design. The 2N4R Tau schematic marks 95 PTMs mapped across 88 positions in human samples [2] and highlights the pS202/pT205/pS208, pT217 and pT231 regions investigated here. The Aβ N-terminal sequence indicates processed and cyclized pyroglutamate at residue 3 (N3pG-Aβ), together with other reported N-terminal modifications. Marks summarize possible modifications and do not imply that all coexist in one molecule. (B) Conceptual uses of compact modification-selective readers for neuronal mapping, biochemical analysis of brain material and prospective reader–E3 or reader–protease effector coupling. Schematics depict potential applications, not established therapeutic outcomes [32]. (C) Two-component design concept: Logos-derived peptide contacts [40] combined with explicit modified-residue motifs and motif-constrained computational generation [42–44].

We reasoned that these challenges could be addressed by designing compact, genetically encoded protein readers that deliberately cooperatively recognize both modification chemistry and the surrounding amino-acid sequence. Unlike phosphotyrosine, phosphoserine and phosphothreonine lack an aromatic ring, placing the phosphate closer to the peptide backbone and limiting hydrophobic anchoring. High-affinity, sequence-specific recognition therefore likely requires both more precise phosphate coordination and cooperative interactions with more surrounding residues. N-terminal pyroglutamate presents a distinct challenge, requiring recognition of a compact cyclic lactam within a short, largely polar sequence. We hypothesized that combining specified sequence-directed peptide recognition [40] with explicit atomic constraints on modified chemical groups [42–44] could address these challenges (Fig. 1C). By establishing complementary interactions with the surrounding peptide and designing a chemically tailored pocket for the modification, both recognition components could be optimized cooperatively. Extending these interfaces linearly toward neighboring residues could further enable tolerance or exclusion of additional PTMs to be programmed prospectively, rather than discovered through empirical characterization. More broadly, we reasoned that these chemically informed recognition principles could be transferable across generative models. Such readers could enable molecularly resolved imaging, biochemical analysis, and ultimately selective perturbation to distinguish disease-driving molecular states from compensatory or incidental changes.

## Results

### Design strategy for chemically defined Tau and Aβ epitopes

We targeted three AD-associated Tau phosphorylation regions—pS202/pT205/pS208, pT217 and pT231—and N-terminal pyroglutamate-modified Aβ (N3pG-Aβ), spanning distinct modification chemistries, peptide sequences and available binding interactions (Fig. 1A). To design readers against these chemically diverse epitopes, we developed a design strategy combining sequence-directed peptide recognition with explicit atomic constraints on the modified chemical groups (Fig. 1C). Rather than generating the entire PTM-binding interface simultaneously, we first established favorable interactions with the surrounding unmodified peptide sequence and subsequently constructed a complementary pocket around the target modification.

We first used Logos [40] to generate candidate binding interfaces against unmodified peptide segments of 6–12 residues. Designs passing AlphaFold 3 (AF3) [45] complex prediction filters were structurally clustered, and the ten highest-scoring, structurally distinct complexes were selected as starting templates for each target. From these complexes, we extracted interface motifs contacting the flanking peptide residues, excluding contacts to the modification site and residues at positions ±2 relative to it. These sequence-recognition motifs were then used as structural constraints for RFdiffusion3 or RFdiffusion4 [43,44]. The target PTM was reintroduced into the modeled complexes using AF3, and its local geometry was allowed to vary during subsequent motif-constrained diffusion. This enabled the generation of a

PTM-centered binding domain with a chemically tailored pocket for phosphoserine, phosphothreonine or pyroglutamate, while preserving the pre-established interactions with the surrounding peptide sequence. The resulting complexes underwent two successive rounds of ProteinMPNN sequence design [39] and AF3 structure prediction and filtering [45]. Candidate readers were selected for predicted interface complementarity to both the modified chemical group and the flanking peptide residues (see Methods). We also adapted RFD4 reward-guided search and filtering to favor the same cooperative chemical- and sequence-recognition features (see Methods). This modular strategy generates designs which integrate chemically precise PTM recognition with extensive sequence-specific contacts, providing a framework for engineering readers against diverse modification chemistries and local sequence contexts.

### De novo readers recognize chemically diverse Tau and Aβ epitopes

We first sought to design binders to four modified peptide epitopes: Tau pT231 (KKVAVVR[pT]PPKSPSS), pT217 (SRTPSLP[pT]PPTRE), pS202/pT205/pS208 (GYSSPG[pS]PG[pT]PG[pS]R), and N3pG-Aβ ([pE]FRHDS). These targets present different design challenges. The triply phosphorylated Tau segment is highly polar, proline-rich and contains few conventional hydrophobic anchoring side chains, whereas N3pG-Aβ provides only a six-residue, largely polar N-terminal recognition context. Both have limited contact surface available to sequence-directed peptide-binding designs [40–41].

For each target, 192 designs were expressed in *E. coli*, purified by nickel-affinity chromatography, and screened by biolayer interferometry (BLI) at a reader concentration of 1 μM against modified and corresponding unmodified biotinylated peptides. Modification-selective hit rates varied across targets: 12/192 (6.25%) for Tau pT231, 7/192 (3.65%) for pT217, 1/192 (0.52%) for pS202/pT205/pS208, and 31/192 (16.1%) for N3pG-Aβ, yielding 51 selective hits among 768 designs (6.64%; Fig. 2A–D). Surface plasmon resonance (SPR) confirmed preferential recognition of modified peptides by representative readers, with apparent dissociation constants of Kd as 1 nM, 83 nM, 121 nM, and 42 nM, respectively. Recognition extended to hyperphosphorylated full-length human Tau and N3pG-Aβ, with minimal binding to corresponding unmodified proteins (Fig. 2A–D).

**Figure 2.**
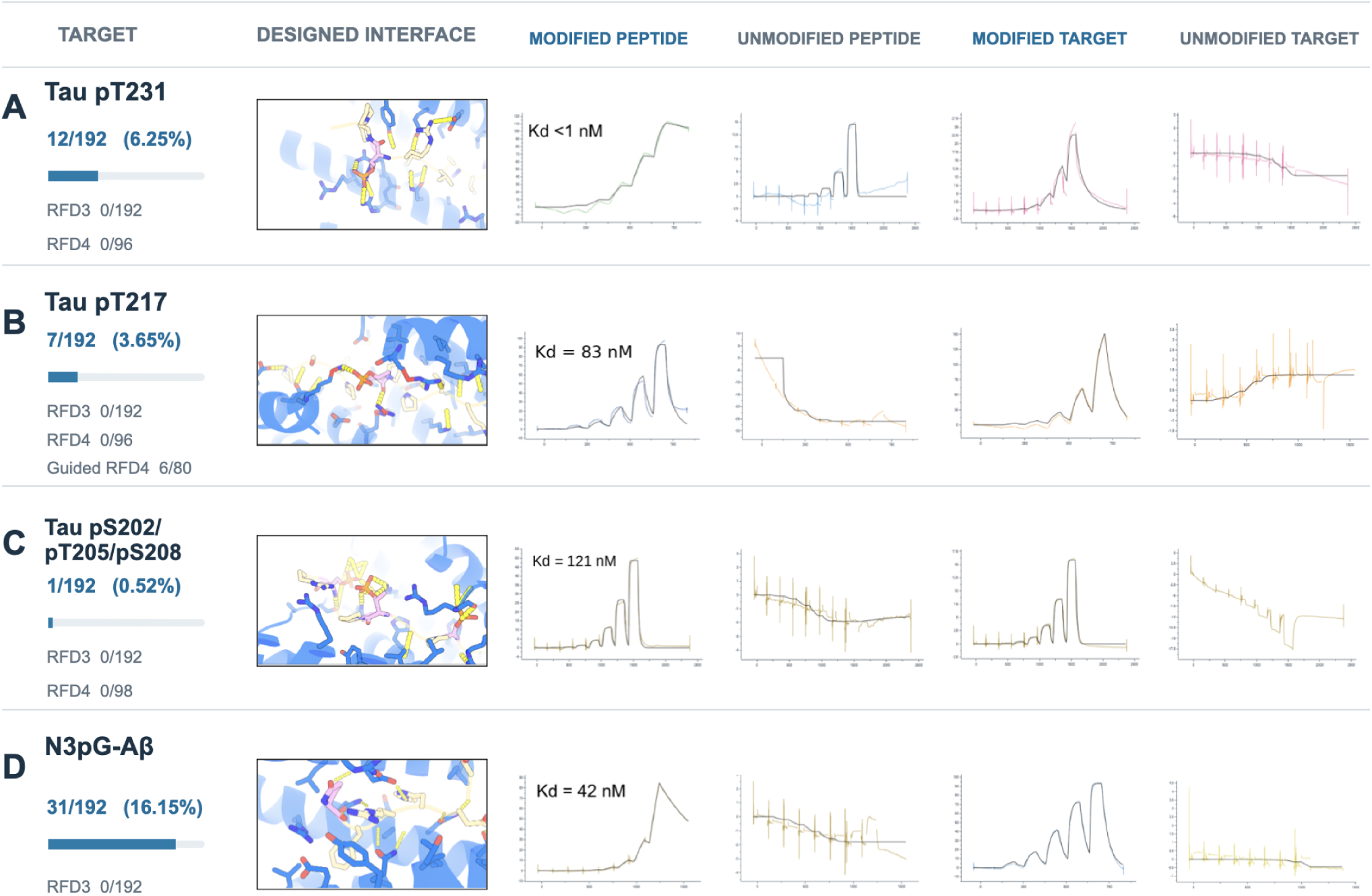
Designed readers recognize modified Tau and Aβ in biochemical assays. (A–D) Representative computational reader–epitope models and binding-response traces for Tau pT231 (A; 12/192 modification-selective hits, 6.25%), Tau pT217 (B; 7/192, 3.65%), Tau pS202/pT205/pS208 (C; 1/192, 0.52%) and N3pG-Aβ (D; 31/192, 16.15%). Each row shows the modeled interface, modified and unmodified peptide interactions, and corresponding longer-target preparations. Reported representative modified-peptide apparent dissociation constants are <1 nM, 83 nM, 121 nM and 42 nM, respectively. The hit fractions represent initial screening, whereas affinities correspond to the selected readers. Binding responses retain their original acquisition scales and cannot be compared quantitatively across panels without normalization. A separate matched-budget RFD3-only campaign is summarized in the lower benchmark panel (0/768 modification-selective hits).

Earlier phosphotyrosine-binder design efforts reported initial success rates often below 0.1%, although some differences in targets, design strategies, and hit criteria preclude direct comparison [42]. To assess the importance of our specificity-guided design strategy, we performed independent baseline campaigns using RFdiffusion3 (RFD3) and RFdiffusion4 (RFD4) without the combined chemical- and sequence-recognition constraints. Baseline RFD3 generated no modification-selective hits experimentally across the same four targets (0/768), while baseline RFD4 likewise yielded no selective hits for Tau pT231 (0/96), pT217 (0/96), or pS202/pT205/pS208 (0/98) experimentally at 1 μM SPR screening condition. Overall, none of the 1,058 baseline designs exhibited modification-selective binding under our screening conditions, compared with 51/768 (6.64%) using our approach. However, incorporating our specificity-guided criteria into RFD4 reward-guided search and filtering recovered 6 selective pT217 readers among 80 designs (7.5%), compared with 0/96 from the baseline RFD4 campaign, all on the same backbone (see Methods). Although these RFD4 readers exhibited weaker affinities, in the high-nanomolar to low-micromolar range, their successful recovery in a smaller tested batch supports the transferability of our recognition principles to a distinct generative framework. Together, these results establish selective recognition across distinct phosphoserine/phosphothreonine and pyroglutamate chemistries and highlight the importance of explicitly guiding cooperative chemical and sequence recognition to achieve modification-selective binding across generative models.

### Designed readers distinguish PTM chemistry and neighboring phosphorylation states

We next evaluated the ability of designed readers to discriminate their intended targets from unrelated modified peptides and chemically similar epitopes. Representative readers were screened at 1 μM against a panel of seven short peptides comprising the three Tau phospho-epitopes, TDP-43 pS409/pS410, α-synuclein pS129, N3pG-Aβ, and gonadotropin-releasing hormone (GnRH), which also bears an N-terminal pyroglutamate (Fig. 3A). Each reader showed a strong response to its intended target, with little or no detectable binding to most noncognate peptides under the tested conditions. Some cross-reactivity was nevertheless observed: the N3pG-Aβ reader also recognized GnRH, albeit with a substantially weaker response. The N3pG-Aβ reader was designed against a minimal six-residue epitope, suggesting that incorporating additional flanking sequence context could further improve specificity. Together, these results demonstrate preferential recognition of the designed epitopes.

**Figure 3.**
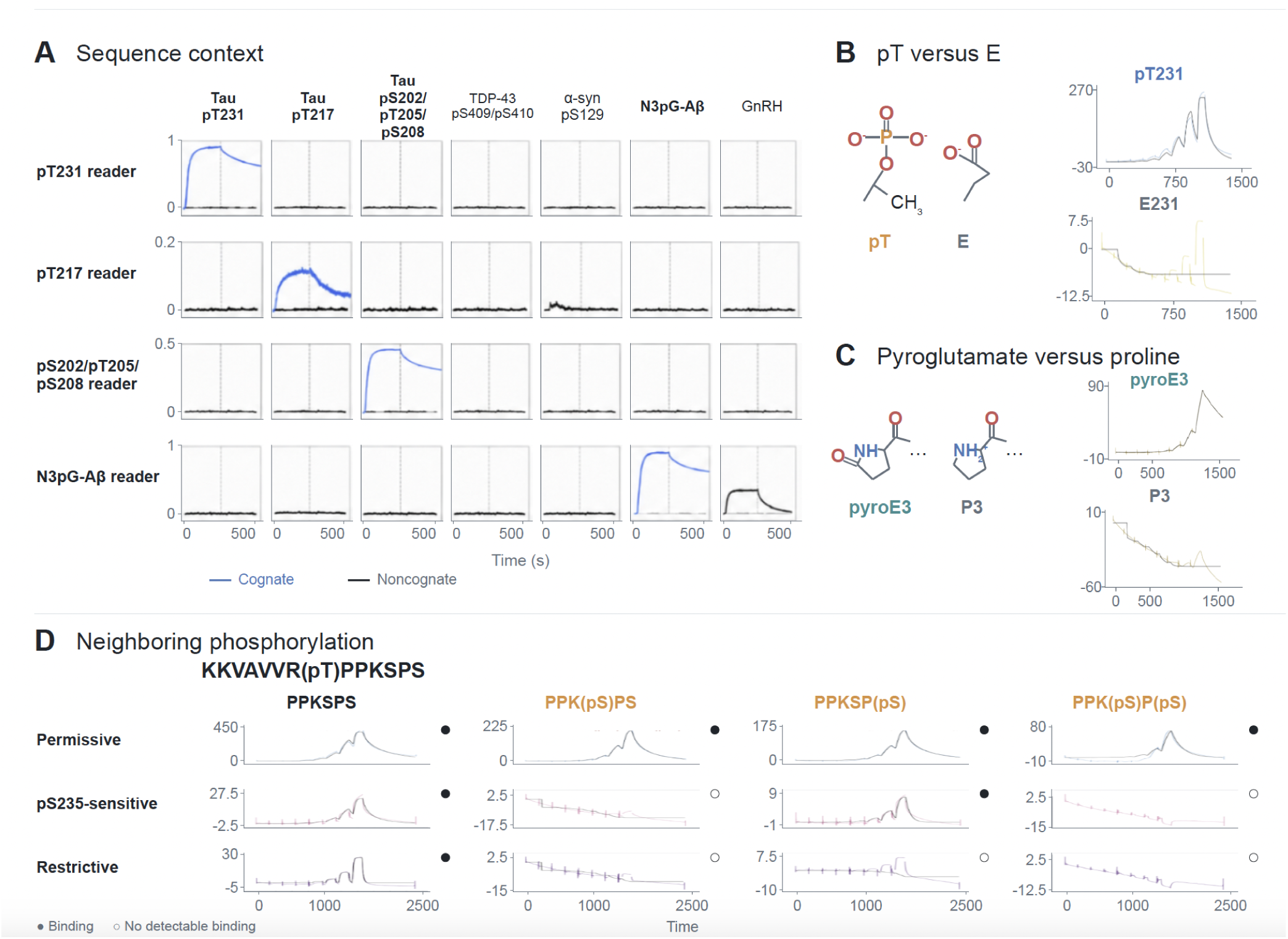
Sequence context, authentic modification chemistry and neighboring phosphorylation determine reader specificity. (A) Four designed reader classes screened against seven peptides: three Tau phospho-epitopes, TDP-43 pS409/pS410, α-synuclein pS129, N3pG-Aβ and GnRH. Blue marks cognate signals; black marks noncognate responses, including pT217-reader cross-reactivity with α-synuclein pS129 and N3pG-Aβ-reader cross-reactivity with GnRH. Response scales vary by row; amplitudes are not affinity ratios. (B) Comparison of binding to authentic Tau pT231 and its E231 substitution. The diagrams illustrate that a glutamate carboxylate does not reproduce the geometry and charge distribution of phosphothreonine [29]. Representative responses are substantially weaker for E231 under the conditions shown. (C) Comparison of N-terminal pyroglutamate at Aβ residue 3 (pE3) with a proline substitution (P3). Although both residues are cyclic, pyroglutamate contains a lactam carbonyl absent from proline; binding responses preferentially recognize the authentic pE3 target. (D) AF3-selected pT231 reader variants [45] tested against peptides with pT231 alone or neighboring pS235 and/or pS237. The three binding classes are permissive, pS235-sensitive and restrictive. Filled/open circles indicate qualitative binding/no detectable binding on panel-specific scales.

We further compared recognition of the authentic modification chemistry with substitution-based mimics. A pT231 reader recognized the phosphorylated Tau peptide but showed little detectable response to a T231E phosphomimetic (Fig. 3B). Glutamate can introduce a negatively charged side chain, but it does not reproduce the geometry or charge distribution of phosphothreonine [29]. Similarly, an N3pG-Aβ reader strongly preferentially recognized N-terminal pyroglutamate over a matched proline substitution (Fig. 3C). Both residues are cyclic, but pyroglutamate contains a lactam carbonyl absent from proline. Thus, similar overall charge or ring shape was not sufficient for binding under these assay conditions, highlighting the specificity of the designs.

Tau molecules carrying the same primary phosphosite can differ in neighboring modifications, contributing to molecular heterogeneity and potentially influencing recognition by phosphosite-specific reagents [2,4,7,24,27]. The T231/S235 region is particularly relevant, as S235 phosphorylation can prime T231 phosphorylation, and modifications at these sites regulate Tau–microtubule interactions [9–10]. We therefore asked whether recognition of neighboring phosphorylation could be deliberately programmed through reader interface redesign. Starting from pT231-selective readers, we extended their binding interfaces toward S235 and S237 through three iterative rounds of ProteinMPNN sequence design and AlphaFold 3 (AF3) structure prediction, selecting variants predicted to differ in their accommodation of neighboring phosphate groups [45]. Three redesigned readers were tested against four matched pT231 peptides containing neither, either, or both additional serine phosphorylations (Fig. 3D). The resulting recognition profiles closely matched the intended designs: one reader tolerated all four phosphorylation contexts; a second tolerated pS237 but not pS235; and a third recognized pT231 only when both neighboring serines were unmodified. These results demonstrate that sensitivity to neighboring PTMs can be tuned through local interface engineering, enabling readers that distinguish modification contexts sharing the same primary phosphosite. Given the heterogeneity and functional consequences of Tau phosphorylation in AD [2,7–8,11,17], such reagents could help resolve molecular populations otherwise grouped together by single-site measurements. Whether these specific S235/S237 phosphorylation combinations define distinct disease-associated populations remains to be established in patient samples.

### Structural validation and interface tuning of PTM readers

To evaluate structural accuracy, we determined the X-ray crystal structure of a reader complexed with the six-residue N3pG-Aβ epitope, (pE)FRHDS, at 2.0 Å resolution (Fig. 4A). Alignment of design and crystal complex showed an interface-backbone root-mean-square deviation (RMSD) of 0.52 Å across 120 matched backbone atoms and an interface side-chain RMSD of 1.64 Å. The N-terminal pyroglutamate is accommodated in a compact pocket, with its peptide-backbone carbonyl positioned 2.82 Å from His43 NE2. The adjacent Phe4 side chain packs against Leu47, Leu51 and Leu97, while Asn94 forms bidentate hydrogen-bonding contacts with the Phe4 backbone amide and carbonyl (3.02 and 3.13 Å). A second bidentate interaction between Asn44 and the Arg5 backbone (2.93 and 2.88 Å), together with His6–Asp91 and Asp7–Ser83 polar contacts, further stabilizes the peptide-binding geometry. These structurally conserved asparagine-mediated backbone interactions extend the recognition principles established in our previous IDR-binder designs [40] to a chemically modified epitope. The close structural agreement highlights the accuracy with which our design approach simultaneously specifies PTM-recognition geometry and complementary interactions with the surrounding substrate sequence, even within a minimal six-residue epitope.

**Figure 4.**
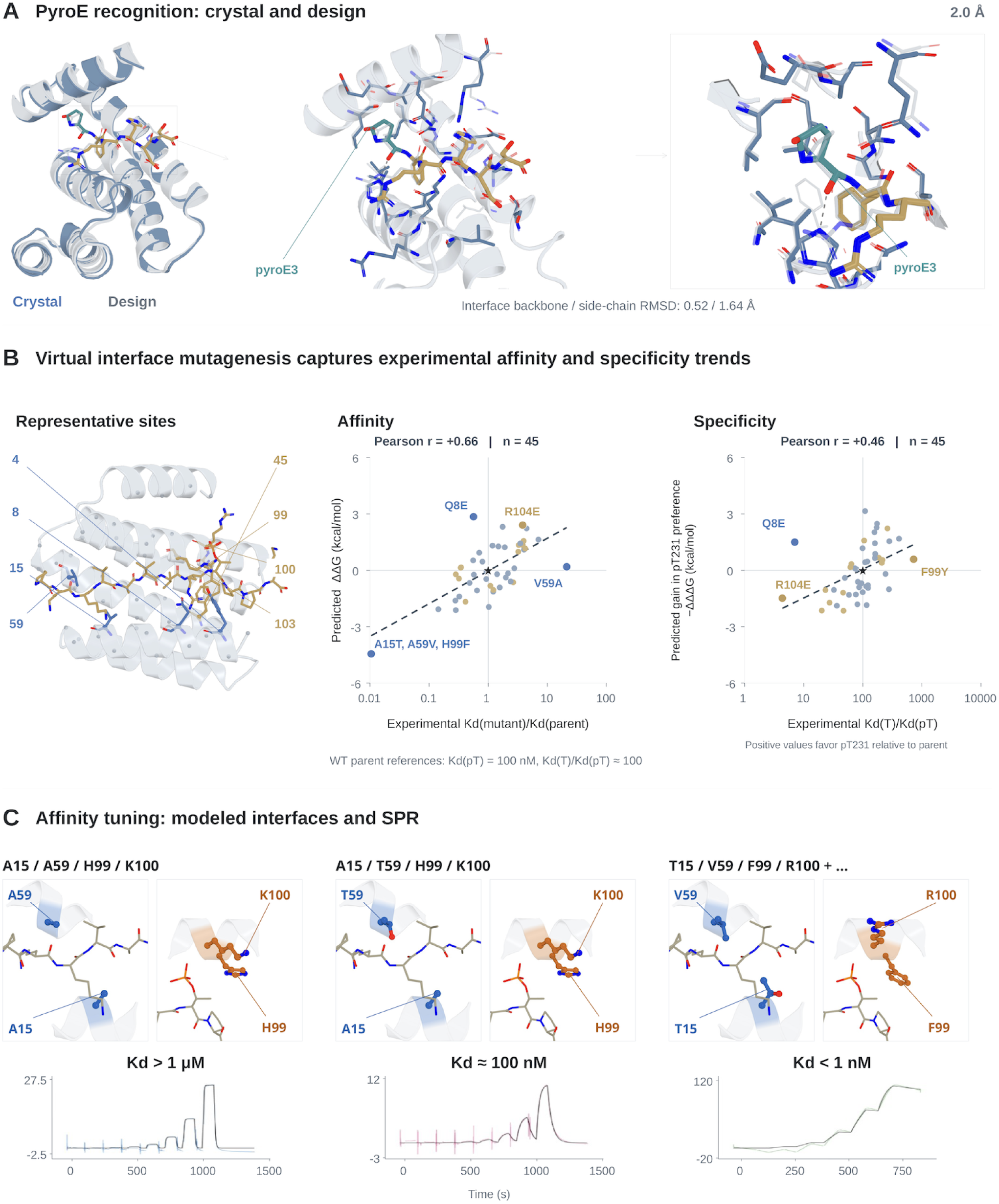
Structural validation and prospective tuning of reader interfaces. (A) X-ray crystal structure (2.0 Å) of an N3pG-Aβ reader bound to (pE)FRHDS compared with its design model. After binder alignment, interface-backbone RMSD is 0.52 Å over 120 matched backbone atoms, and interface side-chain RMSD is 1.64 Å. (B) Modeled pT231-reader contacts and comparison of predicted energetic changes with experimental binding ratios for 45 variants across 8 interface positions. Blue highlights sequence contacts and yellow highlights phosphate-region contacts. The affinity plot compares predicted ΔΔG with Kd(pT, mutant)/Kd(pT, parent). The specificity plot compares predicted gain in pT231 preference (−ΔΔΔG) with Kd(T)/Kd(pT); positive predicted values indicate increased preference for pT231 relative to the parental reference. Dashed lines indicate linear fits against log10-transformed experimental ratios, with Pearson r ≈ 0.66 and 0.46 for affinity and specificity, respectively. Binder numbering begins at residue 1. (C) Paired close-ups of modeled pT231-reader interfaces and representative SPR measurements. Blue highlights side chains at positions 15/59, and orange highlights side chains at positions 99/100. The displayed configurations are A15/A59/H99/K100, A15/T59/H99/K100, and T15/V59/F99/R100, shown alongside representative apparent affinities of >1 μM, approximately 100 nM, and <1 nM, respectively. Matched views facilitate comparison across configurations.

The pT231-reader interfaces contain hydrophobic contacts with the flanking Tau sequence, including at reader positions 15 and 59, alongside interactions near the phosphorylated residue (Fig. 4B). R104 neighbors a terminal phosphate oxygen, whereas R103 lies near the threonine OG1 atom shared by phosphorylated and unmodified threonine, suggesting distinct contributions to affinity and chemical discrimination. To systematically investigate these relationships, we analyzed a virtual library comprising 300 single-, double-, and triple-substitution variants across 8 interface positions and compared predicted energetic changes with experimentally measured binding properties for a subset of 45 variants (Fig. 4B). Computed binding-energy changes correlated with experimental changes in pT231 affinity (Pearson r ≈ 0.66), while predicted increases in pT231 preference correlated with experimental discrimination between pT231 and T231 (r ≈ 0.46). Changes at phosphate-distal positions modulated affinity without proportionate effects on modification specificity, whereas substitutions affecting phosphate coordination, such as R104E, altered the preference for phosphorylated Tau. These results suggest that coupling energy calculations with experimental screens could provide a route to generating reagents with ultrahigh affinity and specificity.

To further illustrate the extent of affinity tuning achievable through interface redesign, we compared representative pT231-reader variants (Fig. 4C). SPR measurements revealed apparent affinities spanning more than three orders of magnitude, from >1 μM for the A15/A59 configuration, through approximately 100 nM for A15/T59, to <1 nM for a multiply substituted interface incorporating T15, V59, F99 and R100. Together with the 2.0 Å N3pG-Aβ reader crystal structure, which closely matches its computational design (interface backbone and side-chain RMSDs of 0.52 and 1.64 Å, respectively; Fig. 4A), these results further demonstrate the structural accuracy, prospective mutational prediction, and extensive affinity tunability of our approach. Such control could enable optimization of readers for distinct biochemical detection, imaging, and cellular applications.

### Designed readers detect AD-associated pathology in neuronal cultures and human brain samples

We next investigated whether readers designed against defined peptide modifications could recognize disease-associated molecular species in complex biological samples. We stained human 3D neuronal cultures derived from AD-like (H10) and control (G10) lines with biotinylated readers targeting N3pG–Aβ (A12), Tau pT231 (A10), and Tau pT217 (AA5), using pathology-associated antibodies as references (Fig. 5A). All three readers produced localized staining that overlapped with pathological antibody signals, with higher reader–antibody colocalization in AD than control cultures. A12 and AA5 also showed substantially increased staining intensity in AD cultures, whereas A10 exhibited more modest differences in overall fluorescence despite its increased colocalization with pathological staining. These results demonstrate that readers designed against individual modified peptides can recognize AD-associated material in human neuronal cultures, with selective staining patterns across the three targets.

**Figure 5.**
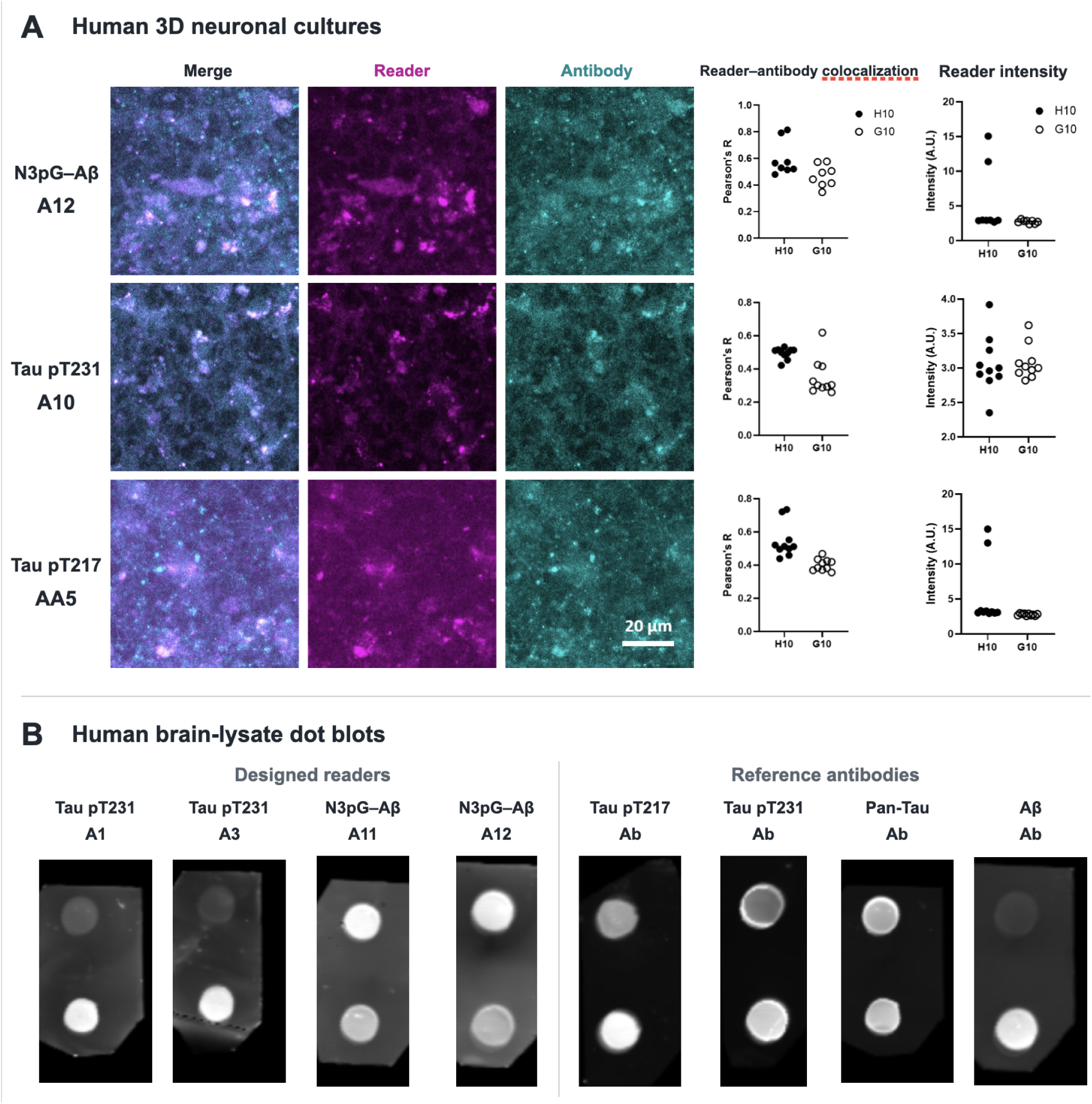
Designed readers detect AD-associated molecular signals in human neuronal cultures and brain lysates. **(A)** Representative fluorescent images of human 3D neuronal cultures stained with biotinylated readers targeting N3pG–Aβ (A12), Tau pT231 (A10), or Tau pT217 (AA5), together with pathology-associated reference antibodies. Reader fluorescence is shown in magenta and antibody fluorescence in cyan; merged images are shown on the left. Quantification compares AD-derived H10 AD-like cultures (filled symbols) with healthy control G10 cultures (open symbols) for reader–antibody colocalization (Pearson’s r) and reader fluorescence intensity (arbitrary units, a.u.). Each point represents one designed reader. Scale bar, 20 μm. **(B)** Dot blots of human brain lysates, probed with Tau pT231 readers A1 and A3, N3pG–Aβ readers A11 and A12, and reference antibodies recognizing Tau pT217, Tau pT231, pan-Tau, or pan-Aβ. Paired lysate spots are shown for each reagent. The pT231 reader blots display an AD-associated signal difference relative to control specimens, whereas pan-Tau staining is comparable across the samples.

We next examined whether designed readers could detect AD-associated Tau species in soluble human brain lysates (Fig. 5B). Such preparations provide an opportunity to investigate non-tangle-associated Tau species, including potentially soluble forms relevant to earlier disease detection. The pT231 readers A1 and A3 preferentially stained AD patient lysates compared with healthy controls, whereas a pan-Tau antibody showed comparable staining across the same samples. Notably, both designed pT231 readers showed clearer discrimination between AD and control lysates than the conventional pT231 antibody, suggesting highly sensitive preferential recognition of disease-associated molecular states.

This contrast suggests that the designed readers recognize disease-associated molecular features rather than simply reflecting differences in total Tau abundance. N3pG–Aβ readers A11 and A12 showed a different staining pattern across the paired brain lysates opposite to that of the pan Aβ reference antibody (Fig. 5B). The molecular basis of this discordance remains unresolved, and the reader signals alone do not establish differences in endogenous N3pG–Aβ abundance. Together, these results extend the activity of designed PTM readers from defined peptide targets to complex neuronal and human brain samples, and highlight their potential for detecting pathological Tau species beyond mature neurofibrillary tangles.

## Discussion

We describe a general computational strategy for designing compact, genetically encoded protein readers that recognize chemically defined post-translational modification (PTM) states, extending de novo protein design beyond recognition of individual peptide sequences or modification sites. By combining sequence-directed binding interfaces with chemically tailored PTM-binding pockets, this approach enables recognition to be specified across three axes: the chemical identity of the modification, the surrounding amino-acid sequence, and the presence or absence of neighboring modifications. The ability to prospectively engineer pT231 readers with distinct tolerance for phosphorylation at S235 and S237 demonstrates that recognition of combinatorial modification states can be programmed through interface design. The close agreement between the designed interface and the 2.0 Å crystal structure of an N3pG-Aβ reader supports the structural accuracy of this approach, while redesign of pT231 interfaces demonstrates that affinity can be tuned over several orders of magnitude. The correlation between virtual mutational energies and experimentally measured changes in affinity and specificity further suggests that chemically informed interface models can guide prospective reader optimization, although improved energetic calibration will be needed for more quantitative prediction. Beyond synthetic peptides, designed readers recognized pathology-associated signals in human 3D neuronal cultures, with spatial correspondence to pathological antibody staining. The pT231 readers preferentially detect soluble AD patient brain lysate Tau, whereas a pan-Tau antibody showed comparable total tau levels in AD and healthy-control samples loaded for this blot. This suggests that designed readers can detect disease-associated molecular features beyond differences in total Tau abundance, potentially including pathological Tau species outside mature neurofibrillary tangles. Together, these results establish a framework for designing molecular readers with predefined recognition properties and applying them to distinguish disease-associated protein states in complex biological samples.

Previous advances in de novo protein design have enabled recognition of intrinsically disordered sequences [40–41] and phosphotyrosine-containing peptides [42], while the accompanying RFD4-Proteina study extends these capabilities to phosphoserine- and phosphothreonine-containing histone epitopes using yeast-display screening [44]. Our results suggest that combining pre-specified peptide contacts with modification-specific interface generation can enable selective recognition by synthetic peptides and purified proteins with higher affinity. The recovery of modification-selective pT217 readers by incorporating chemically informed reward-guided search and filtering into RFD4, but not under the baseline design conditions, provides evidence that these recognition principles can transfer across generative frameworks. Compared with conventional phospho-specific antibodies, whose dependence on neighboring modifications is often discovered retrospectively through empirical characterization [24–28], the ability to prospectively program such preferences offers an additional degree of molecular control. Nevertheless, recognition of defined peptides does not establish proteome-wide specificity or the precise molecular identities of endogenous reader-bound species. A few readers exhibit cross-reactivity with related modified peptides, and their performance varies across biological assays. Differences in epitope accessibility, molecular presentation, sample preparation, and assay readouts may influence recognition independently of intrinsic peptide affinity. In particular, although the human brain-lysate experiments demonstrate discrimination of AD-associated material in the soluble fraction, they do not establish the assembly state or exact proteoform identities of the detected species. The discordant lysate-staining profiles of the N3pG–Aβ readers and pan Aβ reference antibody highlight the distinction between recognition of a defined peptide epitope and quantitative detection of endogenous proteoforms. Differential partitioning or accessibility of pE3-containing species could contribute to this observation, although recognition of different Aβ-derived fragments, non-Aβ material, or sample-dependent assay effects remains possible and is under further investigation. Molecular identification of reader-bound species and analysis of matched soluble and insoluble fractions will be required to distinguish these explanations. Overall, it will be important to determine specificity on a proteome wide level, biochemically characterize reader-bound material, and evaluate binders across independent human samples.

Our approach should enable disease-associated molecular states to be measured and experimentally interrogated with defined specificity. Panels of complementary readers could enable systematic mapping of heterogeneous and combinatorial Tau proteoforms, including soluble pathological species potentially missed by aggregate-focused assays, and help elucidate their relationships to disease progression. With further validation, such tools could complement existing biomarkers. On the therapeutic front, coupling genetically encoded readers to intracellular effectors could enable selective perturbation of defined molecular populations [31–33], extending molecular-state recognition toward causal interrogation.

Ultimately, these capabilities could help distinguish disease-driving, compensatory, and incidental molecular states, establishing predictive relationships between protein sequence, molecular state, and biological function.

## Methods

### Target definitions and peptide reagents

Readers were designed for three human Tau phosphorylation regions (pS202/pT205/pS208, pT217 and pT231) and the N-terminal pyroglutamate-modified Aβ epitope (N3pG-Aβ). Tau residue numbering follows the 441-residue human 2N4R sequence; Aβ numbering follows the mature Aβ peptide. Design inputs consisted of short, initially unmodified peptide segments (typically 6–12 residues) containing the targeted site. The corresponding phosphorylated or pyroglutamyl group was then modeled explicitly for subsequent binder generation. Core peptide sequences and modification states are listed in Table 1; computational design segments and experimentally synthesized peptides are recorded as separate entries when their boundaries differ.

**Table 1.**
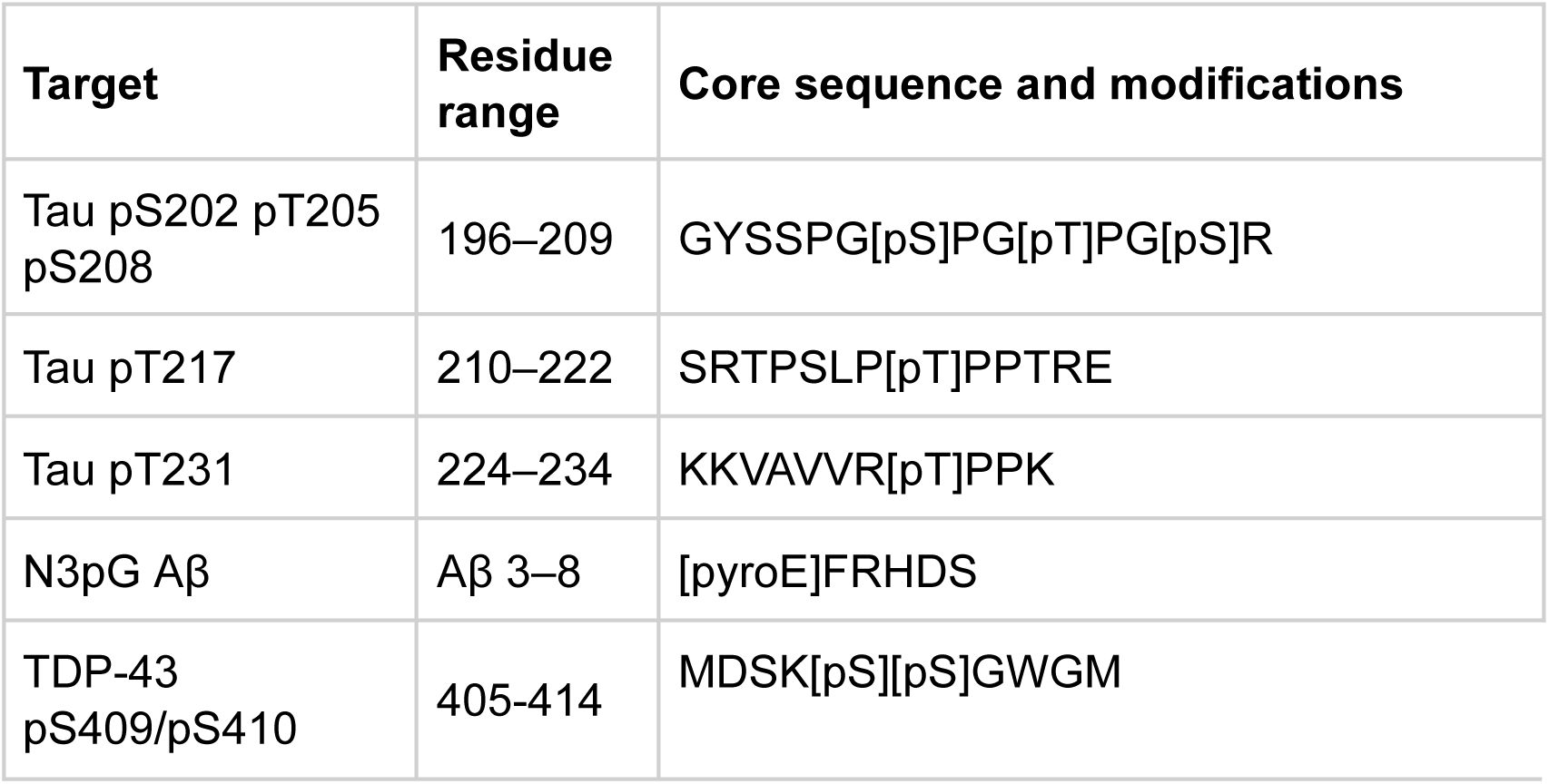
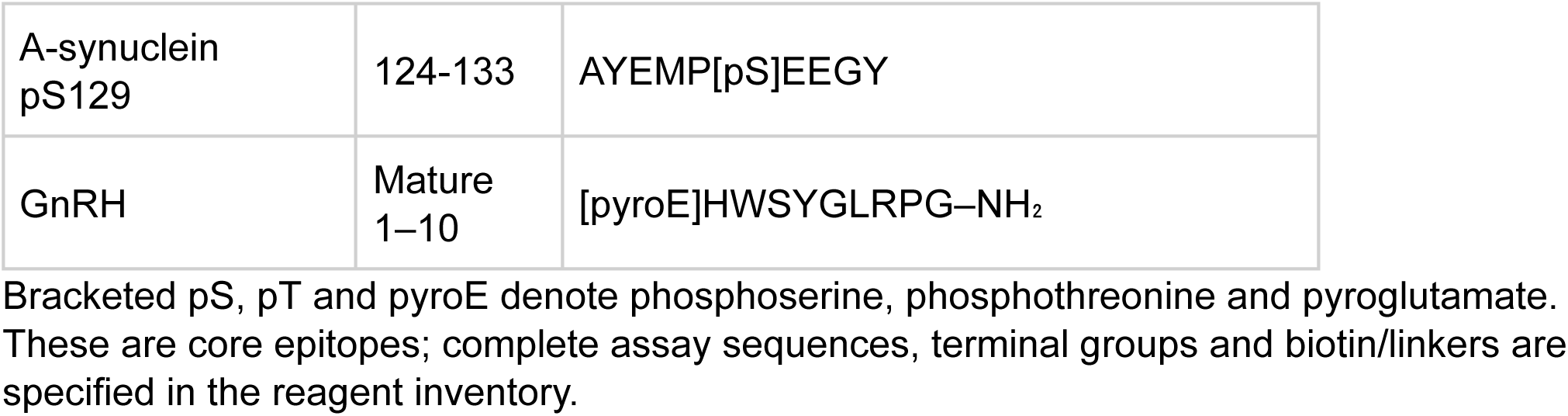
Peptide epitopes used for reader design and specificity characterization.

**Table 2.** Experimentally tested campaign sizes.

| Design arm | Target | Tested designs | Total |
| --- | --- | --- | --- |
| Motif-constrained RFD3 | All four targets | 192 per target | 768 |
| Baseline RFD3 | All four targets | 192 per target | 768 |
| Baseline RFD4 | Tau pT231 | 63 | 96 |
| Baseline RFD4 | Tau pT217 | 96 | 96 |
| Baseline RFD4 | Tau pS202/pT205/pS208 | 98 | 98 |
| Guided RFD4 | Tau pT217 | 80 | 80 |

Counts refer to tested sequences, not total generated candidates. The three baseline RFD4 groups total 290 designs; together with baseline RFD3 they total 1,058. The guided pT217 group is reported separately.

Synthetic target and control peptides were supplied by GenScript or an institutional peptide-synthesis facility and used with the specified residue modifications and terminal chemistries. For each Tau target, a sequence-matched unphosphorylated peptide served as the primary negative control. Additional chemistry controls included a T231E substitution in the pT231 sequence and a proline substitution in place of N-terminal pyroglutamate for the Aβ epitope. Combinatorial-phosphorylation experiments used four versions of the same pT231-containing Tau peptide, differing only in whether S235 and S237 were phosphorylated. Biotinylated peptides were prepared for capture onto streptavidin BLI sensors.

The cross-reactivity panel comprised seven chemically defined targets: Tau pS202/pT205/pS208, Tau pT217, Tau pT231, TDP-43 pS409/pS410, α-synuclein pS129, N3pG-Aβ and gonadotropin-releasing hormone (GnRH), which also carries an N-terminal pyroglutamyl group. Readers were compared against their cognate and noncognate epitopes under matched assay conditions; this panel measures cross-reactivity to the tested peptides, not proteome-wide specificity. All reader sequences and identification codes are linked to the supplementary sequence table.

### Sequence recognition motifs and constrained reader generation

Sequence-directed starting interfaces were generated using Logos [1], following the peptide-template matching approach also used for disordered MYC epitopes [11]. Briefly, each unmodified 6–12-residue target segment was modeled against a library of predesigned peptide–binder architectures representing diverse extended peptide conformations and amino-acid recognition pockets. Designs were evaluated by AlphaFold 3 (AF3) complex prediction [2], filtered for recovery of the intended target–reader interface and structural plausibility, and grouped to reduce near-duplicate binding modes. The ten highest-ranked structurally distinct complexes per target were retained as starting templates.

From each starting complex, we retained reader–peptide interface fragments making favorable interactions with the amino-acid sequence flanking the intended modification. Contacts to the modified residue itself and to peptide residues within ±2 positions of that residue were excluded from the fixed sequence-recognition motif, leaving local space for chemical-specificity design. The target sequence was rebuilt with its authentic phosphorylation or N-terminal pyroglutamate chemistry using AF3. The retained flanking-contact motifs were then provided to RFdiffusion3 (RFD3) or RFD4-Proteina as structural constraints [3,4], while residues and interaction geometry surrounding the modification were allowed to vary. The resulting structures therefore coupled preservation of sequence-directed contacts to de novo sampling of a complementary pocket for the PTM.

Candidate complexes underwent two successive cycles of ProteinMPNN sequence design [5] followed by AF3 complex prediction and filtering. The first cycle generated sequences compatible with the motif-constrained backbones; the second was used to refine sequence–structure agreement and the binding interface while retaining favorable contacts to the authentic modified group and to the surrounding peptide. Designs were ranked for a recovered peptide-binding geometry, plausible PTM coordination, folded-reader confidence and interface complementarity. Structural diversity was retained when selecting the 192 sequences tested per target. The computational design manifest associates each expressed sequence with its target epitope, starting Logos motif, RFD3/RFD4 run, AF3 predictions and model identifier. Further computation details and scripts will be released upon journal acceptance.

### Baseline campaigns and RFD4 reward guided generation

To assess the contribution of cooperative chemical- and sequence-recognition design, independent baseline RFD3 and RFD4 campaigns were carried out without the full motif-constrained, PTM-specific selection strategy. The baseline RFD3 arm screened 192 sequences for each of four epitopes (768 in total); baseline RFD4 arms screened 96 pT231, 96 pT217 and 98 triply phosphorylated Tau designs (290 in total). A separate reward-guided RFD4 pT217 arm favored candidate structures with both a PTM-compatible pocket and contacts to the flanking sequence; 80 sequences were tested. The baseline and guided pT217 collections were nonoverlapping (96 and 80 sequences, respectively). Under the reported modification-selectivity classification, no selective hits were found among 1,058 baseline RFD3/RFD4 candidates, whereas the guided RFD4 pT217 arm yielded six selective hits among 80 tested designs (from the same backbone). These comparisons are between tested design collections, not equivalent total generation budgets.

Baseline and guided designs were expressed and screened against the cognate modified peptide and the corresponding unmodified control using the same 1 µM reader-screening format as the main motif-constrained arm (see ’Reader production and labeling’ and ’Biolayer interferometry screening’). Hit rates were calculated per unique tested sequence and the baseline and reward-guided campaigns were analyzed separately. The RFD4 generative framework is described in [4]; for the reward-guided implementation here, chemical-group complementarity and interactions with nearby peptide sequence were considered during design selection. Further computation details and scripts will be released upon journal acceptance.

### Reader production and labeling

Reader coding sequences were synthesized as linear DNA fragments and subcloned for recombinant expression using the high-throughput peptide-binder production workflow described previously [6]. In that workflow, Golden Gate reactions were assembled in 96-well plates using the LM670 expression vector (Addgene 191552; C-terminal hexahistidine tag), and plasmids were transformed into chemically competent Escherichia coli BL21(DE3).

Following approximately 1 h of recovery in SOC medium, cultures were grown overnight in LB containing kanamycin and then inoculated into approximately 1 ml per well of TBII autoinduction medium supplemented with kanamycin, 2 mM MgSO4 and 1× 5052. Expression was performed for 6 h at 37 °C followed by 18 h at 30 °C. Cultures were harvested at 4,000g for 15 min; pellets were lysed for 15 min in 200 µl 1× BugBuster containing DNase (0.01 mg ml−1) and protease inhibitor, then clarified at 4,000g for 10 min. His-tagged readers were captured on Ni-charged magnetic beads (GenScript L00295), washed in 25 mM Tris (pH 8.0), 300 mM NaCl and 30 mM imidazole, and eluted with the same buffer containing 400 mM imidazole [6].

The earlier IDR pipeline analyzed plate-scale nickel eluates directly by BLI at an estimated final reader concentration of approximately 1 µM, and subsequently expressed selected hits at larger scale [6]. For preparative purification in that workflow, 50-ml autoinduction cultures were grown initially at 37 °C (6 h) and then at 22 °C, lysed in Tris/NaCl buffer with lysozyme, DNase, PMSF and protease inhibitors, clarified at 14,000g, and purified with Ni-NTA agarose followed by size-exclusion chromatography on Superdex 75 Increase 10/300 GL in 25 mM Tris-HCl, 150 mM NaCl (pH 8.0). Purified-protein concentration was estimated from A280 using the sequence-derived extinction coefficient. For the present study, follow-up BLI, SPR and crystallography preparations were associated with their individual purification records.

For the neuronal-culture and brain-lysate applications, individual readers were provided in biotinylated form, enabling detection through a biotin-binding reporter rather than through the streptavidin-immobilized peptide format used in initial BLI screening. Reader biotinylation and peptide biotinylation are distinct reactions and are documented separately. A12 (N3pG-Aβ), A10 (Tau pT231) and AA5 (Tau pT217) were used for the representative neuronal fields; A1 and A3 (Tau pT231) and A11 and A12 (N3pG-Aβ) were among the dot-blot probes.

### Biolayer interferometry screening

High-throughput binding screens were conducted by biolayer interferometry (BLI) at a nominal soluble-reader concentration of 1 µM, using the modified peptide and its unmodified sequence-matched control in parallel. Each of the four motif-constrained campaigns contained 192 experimentally tested reader sequences. The established IDR-binder platform used an Octet RED96 (ForteBio) instrument equipped with streptavidin-coated biosensors (Sartorius 18-5019), with assay buffer comprising 1× HBS-EP+ (Cytiva BR100669) supplemented with 0.1% (w/v) BSA [6]. In that protocol, approximately 50 nM biotinylated peptide was captured on each sensor for 50 s, with 60-s baseline periods before/after loading, an association window of 300–1,200 s and a dissociation window of 200–800 s, adjusted to the target and observed kinetics.

Each binding response was compared with the matching control peptide to discriminate modification-dependent recognition from binding to the underlying sequence. In the earlier IDR BLI method, binding of soluble designs to unloaded streptavidin tips was assessed and the unloaded-tip signal was subtracted from the loaded-sensor trace before association/dissociation fitting [6]. For this study, modification-selective hits were assigned at the unique-sequence level after inspection of cognate modified versus unmodified peptide traces, rather than by counting replicate wells as independent hits. Target-specific hit counts were 12/192 for pT231, 7/192 for pT217, 1/192 for pS202/pT205/pS208 and 31/192 for N3pG-Aβ (51/768 overall).

Peptide-panel specificity measurements used a soluble-reader concentration of 1 µM across the seven-peptide panel described above. Each reader was assayed on its cognate chemically modified peptide and on noncognate peptide targets carrying other phosphorylation or pyroglutamate contexts. Further specificity comparisons assessed a pT231 peptide against the T231E phosphomimetic, N3pG-Aβ against an N-terminal proline substitution, and three redesigns against four pT231/pS235/pS237 phosphorylation states.

Responses were interpreted within each assay using matched concentrations and capture formats; signal amplitude was not treated as a direct affinity ratio. Apparent binding or no-binding calls referred to the sensitivity of the assay under the tested conditions.

### Surface plasmon resonance and affinity analysis

Selected readers were further characterized by surface plasmon resonance (SPR) over a soluble-analyte concentration series rather than from single-concentration responses. We acquired kinetic data on a Biacore 8K using five successive injections in a fivefold series (1.6, 8, 40, 200 and 1,000 nM) followed by an extended dissociation phase. In the PTM experiments, apparent affinities were measured for cognate modified peptides and selected longer target preparations, with matched negative-control surfaces or preparations where available.

Kinetic SPR sensorgrams were evaluated after reference-surface and blank-injection correction. A 1:1 Langmuir interaction model with globally fitted association (kon) and dissociation (koff) rate constants is an appropriate primary analysis for traces adequately described by a single binding mode; KD was calculated as koff/kon. Where the observed kinetics did not support a well-determined two-rate-constant fit, estimates were treated as apparent values or reported as bounds rather than overinterpreted. Reported representative PTM-reader affinities include <1 nM (pT231), approximately 83 nM (pT217), 121 nM (triply phosphorylated Tau) and 42 nM (N3pG-Aβ). These values arise from purified-protein measurements and are not directly comparable with yeast-display apparent affinities.

The assay manifest links each affinity to the complete reader sequence, peptide or longer-target preparation, assay orientation, concentration series, processed and raw traces, fitting method and uncertainty estimate. Purified-protein BLI/SPR measurements are distinguished from the yeast-display half-maximal concentrations reported for histone readers in the accompanying RFD4 study [4].

### Binding to longer target preparations

To test recognition outside the minimal design epitopes, selected readers were evaluated against longer target preparations, including hyperphosphorylated full-length human Tau and N-terminal pyroglutamate-containing Aβ, alongside corresponding unmodified or non-pyroglutamyl controls. In contrast to a single synthetic phosphopeptide, hyperphosphorylated Tau is a heterogeneous e.coli protein preparation and modified by kinases and further confirmed by mass spectrometry. It does not necessarily contain a stoichiometric, uniquely defined pT217 or pT231 proteoform. Longer Aβ preparations were handled as a separate class from the six-residue [pyroE]FRHDS design peptide.

The longer-target binding experiments compared modified and control preparations using the same general BLI/SPR binding framework as the short-peptide measurements, with assay-specific capture geometry and analyte concentration recorded for each run. The sample manifest links each trace to the exact protein or peptide preparation, its tag(s), concentration, modification status, reference surface and reader sequence. The inference from these experiments is recognition of a more complex target preparation, not proof of binding to a single defined full-length Tau proteoform.

### Redesign for neighboring phosphorylation

Structure-guided specificity engineering was performed on pT231-selective parent readers to extend the peptide-contacting interface toward S235 and S237. The candidate complexes were modeled with the authentic pT231 group retained and with additional phosphorylated or unmodified side chains at the neighboring serines. Three iterative rounds of ProteinMPNN sequence design [5] and AF3 complex prediction [2] were used to explore local interface variants: the intended design objectives were (i) accommodation of either neighboring phosphorylation, (ii) tolerance of pS237 but exclusion of pS235 and (iii) recognition only when both S235 and S237 were unmodified. Candidate sequences were selected according to predicted local packing, absence or presence of phosphate-compatible polar groups and retention of the original pT231 recognition mode.

The selected three redesigned readers were tested against four peptides based on the same Tau pT231 sequence: pT231 alone, pT231/pS235, pT231/pS237 and pT231/pS235/pS237. The peptides were compared in a matched biochemical format, permitting a direct qualitative assessment of the effect of each added phosphate. The resulting profiles included a broadly permissive reader recognizing all four contexts, a reader tolerant of pS237 but sensitive to pS235, and a restrictive reader preferentially recognizing pT231 without either neighboring modification.

### Crystallography and structural comparison

A complex between a designed N3pG-Aβ reader and the six-residue Aβ(3–8) peptide, N-terminal [pyroE]FRHDS, was solved by X-ray crystallography at 2.0 Å resolution. The cyclic pyroglutamate is part of the peptide reagent, not a substitution-based structural mimic. The resulting structure was used to evaluate the designed lactam pocket and sequence-directed backbone hydrogen-bonding contacts, including those to Phe4 and Arg5 of Aβ. Using the sitting drop vapor diffusion method, crystallization experiments were set up with 200 nL drops (at 1:1, 2:1, 1:2 protein:crystallization solution ratios) in 96-well plates at 20 °C on a Mosquito LCP from SPT Labtech. Crystallization drops were imaged using UVEX microscopes and a UVEX PS-256 from JAN Scientific. Diffraction quality crystals for R11P2D5 formed after approximately one month in 0.05M sodium chloride, 0.02M glycine, and 33% w/v PEG1000. All crystals were harvested directly from the crystallization drops and flash-frozen in liquid nitrogen.

Diffraction intensities were integrated and scaled, and the complex model was obtained by structure solution followed by iterative coordinate refinement and geometric validation. Map inspection and model refinement were used to assess whether the expected N-terminal pyroglutamyl group and adjacent peptide residues were supported by experimental density. Full data-collection and refinement statistics, including resolution limits, completeness, multiplicity, Rwork/Rfree, geometric quality and ligand geometry, should be reported with the deposited model. Diffraction data was collected at the 17-ID-2 (FMX) beamline of the National Synchrotron Light Source II. X-ray intensities and data reduction were evaluated and integrated using XDS (1) and merged/scaled using Pointless/Aimless in the CCP4 program suite (2). Structure determination and refinement starting phases were obtained by molecular replacement using Phaser (3) using the designed model for the structures. Following molecular replacement, the models were improved using phenix.autobuild (4); efforts were made to reduce model bias by setting rebuild-in-place to false, and using simulated annealing and prime-and-switch phasing. Structures were refined in Phenix (4). Model building was performed using COOT (5). The final model was evaluated using MolProbity (6). Data collection and refinement statistics are recorded in Table 2.

**Table 2:** Crystallographic Table.

|  | **R11P2D5** |
| --- | --- |
| **Data collection** |  |
| Space group | P 2<sub>1</sub> |
| Cell dimensions |  |
| *a*, *b*, *c* (Å) | 33.20, 46.32, 38.27 |
| $\alpha$, $\beta$, $\gamma$ (°) | 90, 112, 90 |
| Resolution (Å) | 35.37 - 1.91 (2.02 - 1.91) |
| *R*<sub>sym</sub> or *R*<sub>merge</sub> | 0.195 (2.320) |
| *I* / σ*I* | 4.6 (1.0) |
| Completeness (%) | 99.3 (98.7) |
| Redundancy | 7.1 (7.3) |
| **Refinement** |  |
| Resolution (Å) | 35.37 - 2.0 (2.15 - 2.0) |
| No. reflections | 58958 (11961) |
| *R*<sub>work</sub> / *R*<sub>free</sub> | 0.2546 (0.3734) / 0.2988 (0.4258) |
| No. atoms |  |
| Protein | 1076 |
| Ligand/ion | 7 |
| Water | 15 |
| *B*-factors |  |
| Protein | 54 |
| Ligand/ion | 67 |
| Water | 52 |
| R.m.s. deviations |  |
| Bond lengths (Å) | 0.013 |
| Bond angles (°) | 0.94 |

\*Single Crystal used for each data collection. \*Values in parentheses are for highest-resolution shell.

Structural similarity was evaluated by superposing the computationally designed reader model onto the crystallographic reader coordinates and subsequently comparing the corresponding peptide–reader interface. The reported interface-backbone RMSD is 0.52 Å across 120 matched backbone atoms, and the reported interface side-chain RMSD is 1.64 Å. Interface comparisons were restricted to corresponding atoms rather than indiscriminately including disordered or nonmatching termini; backbone and side-chain deviations were kept distinct.

Example interface contacts in the crystal include His43 near the pyroglutamyl carbonyl, Phe4 packing against reader Leu47/Leu51/Leu97, and conserved Asn44/Asn94 contacts to peptide backbone atoms.

### Virtual interface mutagenesis and variant measurements

Structure-informed virtual mutagenesis was performed to investigate which pT231 reader contacts primarily contribute to affinity and which might confer discrimination between pT231 and the sequence-matched unphosphorylated T231 peptide. The parent reader pt3_01 carries V15, F45, V59, F99, R100, R103 and R104 in reader-domain numbering. The analyzed virtual library in the current manuscript comprises 45 single-, double- and triple-substitution constructs across eight interface positions; all mutations were interpreted relative to their explicitly defined sequence backgrounds. Candidate sites included positions flanking the phosphothreonine pocket and residues contacting the surrounding peptide backbone or hydrophobic side chains.

Mutant complexes were computationally assessed in the pT231 and unmodified T231 target states to separate predicted changes in binding from predicted changes in PTM preference. Each calculated score was compared to its own sequence-matched parental reference, and the difference between modified and unmodified target calculations was treated as a selectivity-related prediction. The resulting scores were compared with experimentally observed affinity changes for the characterized sequence-matched subset rather than interpreted as calibrated experimental ΔΔG. The manuscript reports a Pearson correlation of approximately 0.66 between predicted and measured affinity changes and a correlation of approximately 0.46 for the predicted selectivity gain.

Variants chosen for experimental follow-up were recombinantly expressed, purified and characterized using the biochemical procedures described above. Interface changes around positions 15 and 59 were associated with a wide dynamic range in apparent affinity, from >1 µM for an A15/A59 configuration through approximately 100 nM for A15/T59 to <1 nM for a high-affinity configuration involving T15, V59, F99 and R100. These abbreviated configurations are not complete construct identities, and neither off-site mutations nor parental backgrounds should be inferred from the abbreviated labels.

### Human neuronal culture and reader staining

Reader staining was examined in human 3D ReNcell cultures designated H10 (AD-like) and G10 (control), derived from the previously described clonal neural progenitor culture system [7–9]. Cells were expanded in Matrigel-coated flasks in ReNcell expansion medium (DMEM/F-12 supplemented with 2 µg/mL heparin, 2% B27, 20 ng/mL EGF, 20 ng/mL bFGF, and penicillin/streptomycin/amphotericin B). For 3D culture, cells were resuspended in differentiation medium (expansion medium without EGF or bFGF), mixed with Matrigel to a final concentration of 2 × 10⁶ cells/mL in 10% Matrigel, and plated at 150 µL per well (3 × 10⁵ cells per well) in 96-well plates. Fresh differentiation medium was added the following day and replaced every 3–4 days. Cultures were differentiated for 4 or 6 weeks.

Cultures were fixed with 4% paraformaldehyde (PFA), permeabilized with 0.4% Triton X-100 in TBST, and blocked with 10% normal goat serum (NGS). Cultures were then incubated with biotinylated readers and reference antibodies in blocking buffer overnight at 4 °C. Biotinylated readers included N3pG-Aβ reader A12, Tau pT231 reader A10, and Tau pT217 reader AA5 for the representative main-figure images, with additional readers included in the broader imaging screen. Reader binding was detected using streptavidin–CF568 (1 µg/mL) for 2 h at room temperature. Reference antibodies included pan-Tau D1M9X (Cell Signaling Technology, CST 46687; 1:1,000), β-amyloid D54D2 (CST 8243; 1:1,000), Tau pT231 (CST 71429; 1:1,000), and Tau pT217 (CST 51625; 1:1,000). Antibody binding was visualized using appropriate fluorescent secondary antibodies [confirm conjugates and dilutions].

### Microscopy and image quantification

Fluorescence images were acquired using a Zeiss high-content automated confocal imaging system. Four fields per well were randomly selected by the imaging system, and image analysis was performed using an automated analysis pipeline. Matched H10 and G10 cultures were imaged using identical acquisition parameters. Reader staining intensity was quantified as background-subtracted mean fluorescence intensity. Reader–antibody colocalization was quantified using Pearson’s correlation coefficient, calculated from corresponding pixels or voxels in the two fluorescence channels.

### Human brain specimens and dot-blot assays

De-identified postmortem human brain specimens from patients with Alzheimer’s disease (AD) and nondemented controls were obtained from the Massachusetts Alzheimer’s Disease Research Center (MADRC) brain bank under Massachusetts General Hospital (MGH) institutional ethics approval. Soluble brain lysates had been prepared previously for an independent study in 20 mM HEPES (pH 7.4) supplemented with protease and phosphatase inhibitors, adjusted to a total protein concentration of 3 mg/mL, and stored in small aliquots at −80 °C.

Aliquots of AD and control brain lysates were applied to nitrocellulose membranes as corresponding dot-blot spots and probed separately with designed readers or conventional epito-specific antibodies. The reader panel included Tau pT231 readers A1 and A3 and N3pG-Aβ readers A11 and A12. Reference antibodies recognized Tau pT217, Tau pT231, total/pan-Tau, and Aβ. Biotinylated readers were detected using LI-COR streptavidin 680, whereas primary antibodies were detected using LI-COR anti-rabbit 800 secondary antibodies. Near-infrared image acquisition and analysis followed previously published LI-COR-based dot-blot procedures [12], with reader-specific detection as described above. Images were acquired for each probe condition to enable qualitative comparison of AD and control specimens.

### Replication and statistical analysis

Screening yield was reported as the number of distinct modification-selective designs divided by the number of unique sequences tested (192 per target for the main design arm), so repeat sensor traces do not inflate the denominator. The four target-specific hit rates were 6.25%, 3.65%, 0.52% and 16.15% for pT231, pT217, pS202/pT205/pS208 and N3pG-Aβ, respectively (51/768 or 6.64% overall). The RFD3 and RFD4 baseline and guided cohorts were analyzed separately. Individual SPR sensorgrams and fitted affinities were treated as measurements on biochemical preparations; multiple injections of one sample were technical replicates. Biological replicate units for brain analysis were independent donors rather than repeat dots, and for neuronal imaging were independent cultures/batches rather than multiple microscopic fields.

Apparent KD values were determined from the stated kinetic or equilibrium model and reported with fitting uncertainty where the underlying traces supported it; measurements with unresolved very tight or weak binding were retained as inequalities rather than replaced by arbitrary point estimates. Pearson correlation coefficients for interface-mutagenesis predictions were computed across paired computational and experimental variant measurements (approximately r = 0.66 for affinity-related changes and r = −0.46 for modification-selectivity comparisons in the current results). No inferential statistical significance, error-bar definition or biological sample size should be asserted without the underlying source records. Descriptive brain dot-blot and neuronal results were distinguished from formal donor- or culture-level inference.

## Data and code availability

The full amino-acid sequences of tested readers, cognate peptides and corresponding PTM/unmodified controls will be supplied in supplementary sequence and reagent tables, with links from reader IDs to design configurations, AlphaFold predictions and experimental assays. Where permitted, deposited source data will include raw and processed BLI/SPR traces and fitting files, virtual mutagenesis records, original fluorescence images/analysis files and uncropped brain dot blots. Analysis scripts and model/configuration files are planned for deposition on GitHub at or before journal publication, and coordinates plus diffraction structure factors for the N3pG-Aβ complex will be made available through the Protein Data Bank. Human-donor metadata will be shared only to the extent consistent with consent, ethics determinations and applicable access rules.

## Author contributions

K.W., Y.S., and D.B. conceived the study. K.W. developed the computational design strategy, led reader design and characterization, analyzed the results, and coordinated the project. T.S. and L.C. contributed to a large amount of computational design and filtering strategies, interface optimization, and protein biochemistry characterization. P.C. led biological characterization of designed readers, with contributions from E.B. and T.R.K. to neuronal culture experiments and imaging. D.Y.K. provided expertise and supervision for the human 3D neuronal culture models. A.C. contributed to scientific discussions regarding the design process. N.C. contributed to RFD4-based computational design and analysis. A.K.B., A.K., J.M., and H.N. contributed to X-ray crystallographic characterization of designed reader complexes. X.L. contributed peptide synthesis. M.S.B. contributed scientific discussions on PTM-specific binder design. M.B. and B.C.K. contributed to the full length phosphorylated Tau sample preparations. Y.S. guided disease-relevant target selection and supervised biological validation. D.B. supervised protein design research. K.W. wrote the original manuscript with input from Y.S. and D.B. All authors reviewed and approved the manuscript.

## Acknowledgments

We thank K.A.V. for wet-lab management and experimental coordination; L.G. and P.V. for computational infrastructure and IT support; and J.E. and J.C. for valuable scientific discussions, strategic advice, and guidance on research funding. We are grateful to members of the Baker, Song, Kim, and Kraemer laboratories for helpful discussions and technical assistance. This research used resources 17-ID-2 (FMX) of the National Synchrotron Light Source II, a US Department of Energy (DOE) Office of Science User Facility operated for the DOE Office of Science by Brookhaven National Laboratory under Contract No. DE-SC0012704. The Center for BioMolecular Structure (CBMS) is primarily supported by the National Institutes of Health, National Institute of General Medical Sciences (NIGMS) through a Center Core P30 grant (no. P30GM133893) and by the DOE Office of Biological and Environmental Research (grant no. KP1605010). We also acknowledge the individuals and families whose donated brain tissue made these studies possible, as well as the MADRC brain bank for providing the human samples.

## Data and code availability

The amino acid sequences of the designed readers will be provided in Supplementary Table 1 upon journal acceptance. Structural coordinates and experimental data supporting the conclusions of this study will be available through relevant PDB accession, data repository, and code repository, as applicable. Additional source data and analysis scripts are available from the corresponding authors upon reasonable request upon journal acceptance.

## References

1. Wu, K. et al. Design of intrinsically disordered region binding proteins. Science 389, eadr8063 (2025). 10.1126/science.adr8063

2. Abramson, J. et al. Accurate structure prediction of biomolecular interactions with AlphaFold 3. Nature 630, 493–500 (2024). 10.1038/s41586-024-07487-w

3. Butcher, J. et al. Atomic functional site scaffolding in biopolymers with RFdiffusion3. Science (2026). 10.1126/science.aeh2982

4. Corley, N. et al. Biomolecular design through latent reasoning with RFD4-Proteina. Manuscript (2026).

5. Dauparas, J. et al. Robust deep learning-based protein sequence design using ProteinMPNN. Science 378, 49–56 (2022). 10.1126/science.add2187

6. Liu, C., Wu, K. et al. Diffusing protein binders to intrinsically disordered proteins. Nature 644, 809–817 (2025). 10.1038/s41586-025-09248-9

7. Choi, S. H. et al. A three-dimensional human neural cell culture model of Alzheimer’s disease. Nature 515, 274–278 (2014). 10.1038/nature13800

8. Kim, Y. H. et al. A 3D human neural cell culture system for modeling Alzheimer’s disease. Nature Protocols 10, 985–1006 (2015). 10.1038/nprot.2015.065

9. Kwak, S. S. et al. Amyloid-β42/40 ratio drives tau pathology in 3D human neural cell culture models of Alzheimer’s disease. Nature Communications 11, 1377 (2020). 10.1038/s41467-020-15120-3

10. Dunn, K. W., Kamocka, M. M. & McDonald, J. H. A practical guide to evaluating colocalization in biological microscopy. American Journal of Physiology Cell Physiology 300, C723–C742 (2011). 10.1152/ajpcell.00462.2010

11. McNamara-Bordewick, N. K. et al. MYC targeted de novo binders impair tumor growth. bioRxiv (2026). 10.64898/2026.10.15.758618

12. Brady, S. T. et al. Toxic effects of mutant huntingtin in axons are mediated by its proline-rich domain. Brain 147, 2098–2113 (2024). 10.1093/brain/awad280

## References

1. Braak, H. & Braak, E. Neuropathological stageing of Alzheimer-related changes. Acta Neuropathol. 82, 239–259 (1991). 10.1007/BF00308809

2. Wesseling, H. et al. Tau PTM profiles identify patient heterogeneity and stages of Alzheimer’s disease. Cell 183, 1699–1713.e13 (2020). 10.1016/j.cell.2020.10.029

3. Hanger, D. P. et al. Novel phosphorylation sites in tau from Alzheimer brain support a role for casein kinase 1 in disease pathogenesis. J. Biol. Chem. 282, 23645–23654 (2007). 10.1074/jbc.M703269200

4. Dujardin, S. et al. Tau molecular diversity contributes to clinical heterogeneity in Alzheimer’s disease. Nat. Med. 26, 1256–1263 (2020). 10.1038/s41591-020-0938-9

5. Arakhamia, T. et al. Posttranslational modifications mediate the structural diversity of tauopathy strains. Cell 180, 633–644.e12 (2020). 10.1016/j.cell.2020.01.027

6. Fitzpatrick, A. W. P. et al. Cryo-EM structures of tau filaments from Alzheimer’s disease. Nature 547, 185–190 (2017). 10.1038/nature23002

7. Joly, J. et al. Large-scale single-molecule analysis of tau proteoforms. Nat. Methods 23, 1786–1797 (2026). 10.1038/s41592-026-03188-6

8. Fan, X. et al. A pathological phosphorylation pattern enhances tau cooperativity on microtubules and facilitates tau filament assembly. *Commun*. Biol. (2026). 10.1038/s42003-026-10797-6

9. Sengupta, A. et al. Phosphorylation of tau at both Thr 231 and Ser 262 is required for maximal inhibition of its binding to microtubules. Arch. Biochem. Biophys. 357, 299–309 (1998). 10.1006/abbi.1998.0813

10. Cho, J.-H. & Johnson, G. V. W. Primed phosphorylation of tau at Thr231 by glycogen synthase kinase 3β (GSK3β) plays a critical role in regulating tau’s ability to bind and stabilize microtubules. J. Neurochem. 88, 349–358 (2004). 10.1111/j.1471-4159.2004.02155.x

11. Milà-Alomà, M. et al. Plasma p-tau231 and p-tau217 as state markers of amyloid-β pathology in preclinical Alzheimer’s disease. Nat. Med. 28, 1797–1801 (2022). 10.1038/s41591-022-01925-w

12. Palmqvist, S. et al. Discriminative accuracy of plasma phospho-tau217 for Alzheimer disease vs other neurodegenerative disorders. JAMA 324, 772–781 (2020). 10.1001/jama.2020.12134

13. Ashton, N. J. et al. Diagnostic accuracy of a plasma phosphorylated tau 217 immunoassay for Alzheimer disease pathology. JAMA Neurol. 81, 255–263 (2024). 10.1001/jamaneurol.2023.5319

14. Barthélemy, N. R. et al. Blood plasma phosphorylated-tau isoforms track CNS change in Alzheimer’s disease. J. Exp. Med. 217, e20200861 (2020). 10.1084/jem.20200861

15. Janelidze, S. et al. Plasma P-tau181 in Alzheimer’s disease: relationship to other biomarkers, differential diagnosis, neuropathology and longitudinal progression to Alzheimer’s dementia. Nat. Med. 26, 379–386 (2020). 10.1038/s41591-020-0755-1

16. Thijssen, E. H. et al. Diagnostic value of plasma phosphorylated tau181 in Alzheimer’s disease and frontotemporal lobar degeneration. Nat. Med. 26, 387–397 (2020). 10.1038/s41591-020-0762-2

17. Wojdała, A. L. et al. Immunoassay detection of multiphosphorylated tau proteoforms as cerebrospinal fluid and plasma Alzheimer’s disease biomarkers. Nat. Commun. 16, 214 (2025). 10.1038/s41467-024-54878-8

18. Saido, T. C. et al. Dominant and differential deposition of distinct beta-amyloid peptide species, A beta N3(pE), in senile plaques. *Neuron* **14**, 457-466 (1995). 10.1016/0896-6273(95)90301-1

19. Schilling, S. et al. Glutaminyl cyclase inhibition attenuates pyroglutamate Aβ and Alzheimer’s disease-like pathology. Nat. Med. 14, 1106–1111 (2008). 10.1038/nm.1872

20. Nussbaum, J. M. et al. Prion-like behaviour and tau-dependent cytotoxicity of pyroglutamylated amyloid-β. Nature 485, 651–655 (2012). 10.1038/nature11060

21. DeMattos, R. B. et al. A plaque-specific antibody clears existing β-amyloid plaques in Alzheimer’s disease mice. Neuron 76, 908–920 (2012). 10.1016/j.neuron.2012.10.029

22. Sims, J. R. et al. Donanemab in early symptomatic Alzheimer disease: the TRAILBLAZER-ALZ 2 randomized clinical trial. JAMA 330, 512–527 (2023). 10.1001/jama.2023.13239

23. Goedert, M., Jakes, R. & Vanmechelen, E. Monoclonal antibody AT8 recognises tau protein phosphorylated at both serine 202 and threonine 205. Neurosci. Lett. 189, 167–169 (1995). 10.1016/0304-3940(95)11484-E

24. Malia, T. J. et al. Epitope mapping and structural basis for the recognition of phosphorylated tau by the anti-tau antibody AT8. Proteins 84, 427–434 (2016). 10.1002/prot.24988

25. Triana-Baltzer, G. et al. Development and validation of a high-sensitivity assay for measuring p217+tau in plasma. Alzheimers Dement. (Amst.) 13, e12204 (2021). 10.1002/dad2.12204

26. Amniai, L., Lippens, G. & Landrieu, I. Characterization of the AT180 epitope of phosphorylated Tau protein by a combined nuclear magnetic resonance and fluorescence spectroscopy approach. Biochem. Biophys. Res. Commun. 412, 743–746 (2011). 10.1016/j.bbrc.2011.08.046

27. Ercan, E. et al. A validated antibody panel for the characterization of tau post-translational modifications. Mol. Neurodegener. 12, 87 (2017). 10.1186/s13024-017-0229-1

28. Min, S. et al. Effects of neighboring phosphorylation events on the affinities of pT181-Tau antibodies. Anal. Chem. 95, 18241–18248 (2023). 10.1021/acs.analchem.3c04081

29. Bilbrough, T., Piemontese, E. & Seitz, O. Dissecting the role of protein phosphorylation: a chemical biology toolbox. Chem. Soc. Rev. 51, 5691–5730 (2022). 10.1039/D1CS00991E

30. Chan, A. & Tsourkas, A. Intracellular protein delivery: approaches, challenges, and clinical applications. BME Front. 5, 0035 (2024). 10.34133/bmef.0035

31. Clift, D. et al. A method for the acute and rapid degradation of endogenous proteins. Cell 171, 1692–1706.e18 (2017). 10.1016/j.cell.2017.10.033

32. Benn, J. et al. Aggregate-selective removal of pathological tau by clustering-activated degraders. Science 385, 1009–1016 (2024). 10.1126/science.adp5186

33. Silva, M. C. et al. Targeted degradation of aberrant tau in frontotemporal dementia patient-derived neuronal cell models. eLife 8, e45457 (2019). 10.7554/eLife.45457

34. Banik, S. M. et al. Lysosome-targeting chimaeras for degradation of extracellular proteins. Nature 584, 291–297 (2020). 10.1038/s41586-020-2545-9

35. Cao, L. et al. Design of protein-binding proteins from the target structure alone. Nature 605, 551–560 (2022). 10.1038/s41586-022-04654-9

36. Pacesa, M. et al. One-shot design of functional protein binders with BindCraft. Nature 646, 483–492 (2025). 10.1038/s41586-025-09429-6

37. Watson, J. L. et al. De novo design of protein structure and function with RFdiffusion. Nature 620, 1089–1100 (2023). 10.1038/s41586-023-06415-8

38. Anishchenko, I. et al. De novo protein design by deep network hallucination. Nature 600, 547–552 (2021). 10.1038/s41586-021-04184-w

39. Dauparas, J. et al. Robust deep learning-based protein sequence design using ProteinMPNN. Science 378, 49–56 (2022). 10.1126/science.add2187

40. Wu, K. et al. Design of intrinsically disordered region binding proteins. Science 389, eadr8063 (2025). 10.1126/science.adr8063

41. Liu, C. et al. Diffusing protein binders to intrinsically disordered proteins. Nature 644, 809–817 (2025). 10.1038/s41586-025-09248-9

42. Bauer, M. S. et al. De novo design of phosphotyrosine peptide binders. bioRxiv (2025). 10.1101/2025.09.29.678898

43. Butcher, J. et al. Atomic functional site scaffolding in biopolymers with RFdiffusion3. Science, eaeh2982 (2026). 10.1126/science.aeh2982

58. . Corley, N., et al. Biomolecular design through latent reasoning with RFD4-Proteina. preprint (2026).

45. Abramson, J. et al. Accurate structure prediction of biomolecular interactions with AlphaFold 3. Nature 630, 493–500 (2024). 10.1038/s41586-024-07487-w

46. Choi, S. H. et al. A three-dimensional human neural cell culture model of Alzheimer’s disease. Nature 515, 274–278 (2014). 10.1038/nature13800

47. Kim, Y. H. et al. A 3D human neural cell culture system for modeling Alzheimer’s disease. Nat. Protoc. 10, 985–1006 (2015). 10.1038/nprot.2015.065

48. Park, J. et al. A 3D human triculture system modeling neurodegeneration and neuroinflammation in Alzheimer’s disease. Nat. Neurosci. 21, 941–951 (2018). 10.1038/s41593-018-0175-4

